# Focusing perceptual attention in the past constrains outcome-based learning in the future by adjusting cortico-cortical interactions

**DOI:** 10.1101/2022.01.22.477334

**Authors:** Dmitrii Vasilev, Negar Safaei, Ryo Iwai, Hessam Bahmani, Ioannis S. Zouridis, Masataka Watanabe, Nikos K. Logothetis, Nelson K. Totah

**Affiliations:** Helsinki Institute of Life Science (HILIFE), University of Helsinki, 00014 Helsinki, Finland; Department of Physiology of Cognitive Processes, MPI Biological Cybernetics, 72076 Tübingen, Germany; Faculty of Engineering, The University of Tokyo, Tokyo 111-0033, Japan; Division of Imaging Sci. and Biomed. Eng., University of Manchester, Manchester M13 9PL, UK; Dept. of Physiology of Cognitive Processes, International Center for Primate Brain Research, Center for Excellence in Brain Science and Intelligence Technology, Institute of Neuroscience (CEBSIT, ION), Chinese Academy of Sciences (CAS), 201602 Shanghai, China; Faculty of Pharmacy, University of Helsinki, 00014 Helsinki, Finland

## Abstract

Contemporary neuroscience and psychiatry suggest that attention to decision outcomes guides rule learning by adjusting stimulus-outcome associations. Separately, sensory neurophysiology conceptualizes attention as a ‘filter’ that improves perception. Here, we show that the contemporary view is incomplete by demonstrating an unconventional and novel effect of perceptual attention on subsequent outcome-based rule learning. Moreover, we show for the first time in rodents that, like in primates, this attentional process involves tuning of modality specific cortico-cortical interactions. We designed a novel head-fixed rat-on-a-treadmill apparatus and used it to train rats to discriminate auditory-visual stimuli using one modality and then reduced stimulus discriminability in that modality. We observed perceptual learning suggesting engagement of perceptual attention. Moreover, engaging visual perceptual attention resulted in more saccades and increased frontal-visual cortex EEG Granger causality relative to engaging auditory perceptual attention. We then presented novel and easily discriminable stimuli in both modalities and measured outcome-driven learning in the other modality. Learning was slower after engaging perceptual attention. Our work suggests that a more complete description of learning requires integrating these previously siloed concepts of attention. Moreover, treating impaired set-shifting as a trans-diagnostic symptom may require targeting different neural circuits for perceptual attention or outcome-based attention depending on which type of attention is impaired in each neuro-psychiatric disorder.

## Introduction

Attention is critical for learning outcomes predicted by stimuli in a changing environment ^1,2^. Outcome-related attention has been studied using the attentional set-shifting paradigm, in which the rewarded dimension (e.g., color) of multi-dimensional stimuli (e.g., various colored shapes) must be learned through trial-and-error. When presented with novel stimuli, learning to respond to a previously unrewarded stimulus dimension (shape) is slow ^3–7^, presumably because attention is ‘stuck’ on the previously rewarded dimension (color) and must be shifted to the newly rewarded dimension (shape) ^3,8^.

Attention has also been studied as a sensory filter that improves the ability to discriminate similar stimuli by altering sensory neuron representations ^9–11^. This other conceptualization of attention leads to the intriguing question of whether focusing perceptual attention onto one dimension of the environment could influence shifting attention during subsequent outcome-based learning. Answering this question not only characterizes more precisely the role of attention in learning, but also clarifies which forms of attention could underlie attentional set-shifting impairment in individuals diagnosed with attention deficit hyperactivity disorder, autism, obsessive-compulsive disorder, schizophrenia, or substance use disorder ^12–16^.

We tested the role of perceptual attention and concomitant changes in neuronal representations on subsequent learning in a novel attentional set-shifting paradigm. We trained head-fixed rats in Go/NoGo auditory-visual attentional set-shifting task. Prior to assessing the ability to learn responding to novel stimuli using the previously unrewarded modality, perceptual attention was manipulated by presenting either difficult or easy discriminations in the currently rewarded modality. We observed perceptual learning (i.e., the ability to tell apart similar stimuli improves with task experience) during difficult discriminations, which suggests that perceptual attention was engaged in that task condition ^17–22^. During difficult discriminations, we also observed increased EEG Granger causality magnitude between frontal cortex and sensory cortex neurons tuned to the currently rewarded modality. Critically, we observed that learning to respond to novel and easily discriminable stimuli in the other modality was slower after subjecting rats to difficult discriminations in the previously rewarded modality.

## Results

We studied how engaging perceptual attention onto one sensory modality affects subsequent outcome-based learning in another modality using an auditory-visual attentional set-shifting task for head-fixed rats. The task presented compound auditory-visual stimuli constructed from pure frequency tones and visual drifting gratings (**Figure 1A**). These stimuli were chosen because the relevant feature in each sensory modality (i.e., grating orientation or tone frequency) could be parametrically varied to manipulate stimulus discriminability and engage perceptual attention. Rats were initially trained to associate specific responses (Go response – treadmill running; NoGo response – immobility) with stimuli in one modality (**Figure 1B**). The two stimuli in the irrelevant modality were each presented an equal number of times with the Go stimulus (300 trials/session) and the NoGo stimulus (300 trials/session) and in randomized order. We trained 6 rats to perform auditory discrimination and 9 rats to perform visual discrimination (N = 15 rats). Stimuli in both modalities were easily discriminable based on prior visual and auditory psychophysics experiments in rodents ^23–27^.

**Figure 1.**
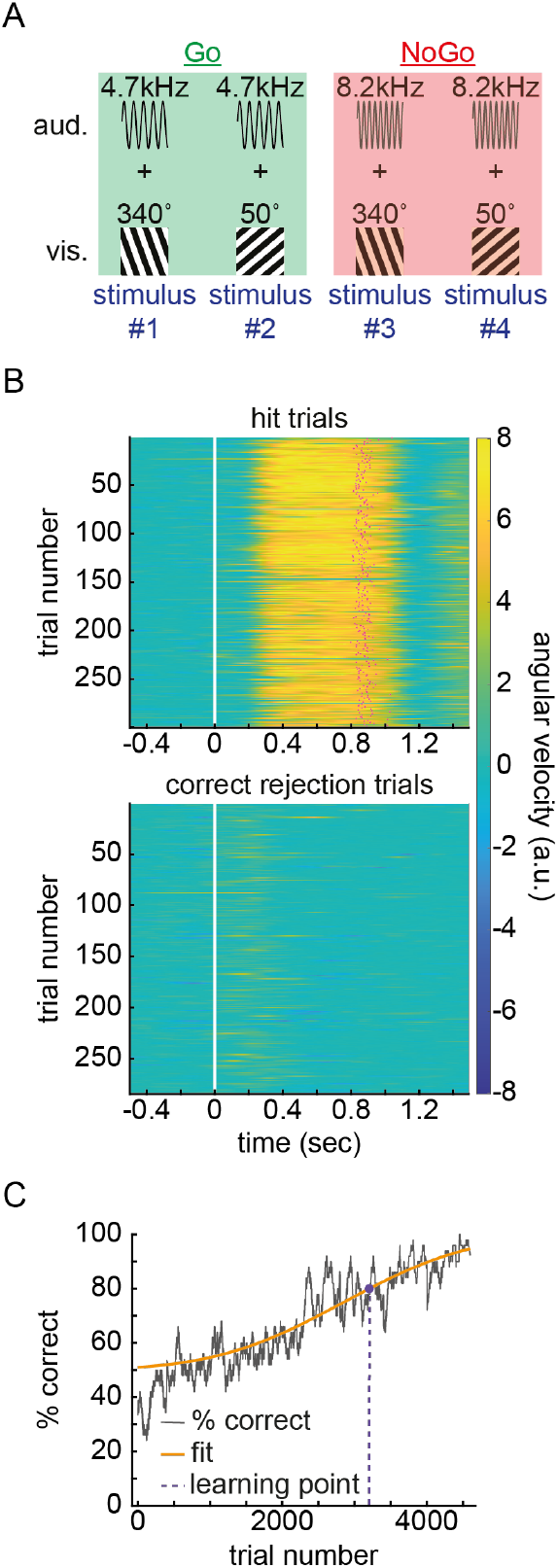
The auditory-visual attentional set-shifting task and the method for defining similar levels of stimulus discriminability across rats. **(A)** An example stimulus set (Go: 4.7 kHz tone; NoGo: 8.2 kHz tone randomly coupled with a 50° or 340° visual drifting grating). **(B)** Rats were trained to remain immobile for 0.5 sec prior to sensory stimulus onset. After stimulus presentation, rats could choose to run or remain immobile. Heatmaps show angular velocity across trials during one session (rotary encoder voltage change/sec scaled by 10^3^). Positive velocity (yellow) indicates forward movement and negative velocity (blue) indicates backwards movement. Near-zero velocity indicates immobility. Backward movement was rare. The magenta dots indicate the time of response threshold crossing on hit trials. The white line indicates stimulus onset. **(C)** The learning curve from an IDS of an example rat. The % correct responses (50 trial bins) was fit with a Sigmoid function. The trial bin at which the fitted line reached 80% is the learning timepoint.

The experiment tested the hypothesis that engaging perceptual attention onto one modality would affect learning the responses associated with novel stimuli in the other modality. The experiment had three steps repeated once. First, we obtained a learning baseline during an intra-dimensional shift (IDS). In the IDS, the ability to learn the responses associated with novel stimuli was assessed when the relevant modality was unchanged. Stimuli in both modalities were easily discriminable. A new session (600 trials) was completed each day until learning finished (i.e., the mean performance for an entire session was >85%). The baseline learning timepoint was defined as the trial at which 80% correct was reached on a Sigmoid function fit to the binned % correct performance calculated in 50 trial bins (**Figure 1C**).

After measuring baseline learning during an IDS, we manipulated perceptual attention onto the relevant modality of the learned stimuli. During two consecutive attention manipulation sessions, perceptual attention was manipulated by requiring either easy or difficult discriminations in the currently relevant modality. Seven rats were presented with the easily discriminable stimuli already learned during the IDS. Eight rats were presented difficult to discriminate stimuli in the relevant modality by changing the NoGo stimulus feature to have greater similarity with the Go stimulus, without modifying the Go stimulus or the stimuli in the irrelevant modality. In other words, stimuli identical to those learned during the IDS were presented, with the exception that the NoGo stimulus was less discriminable from the Go stimulus. Discrimination difficulty was set to a common level across rats using a psychophysics staircase procedure designed to estimate the NoGo stimulus orientation (when vision was relevant) or frequency (when audition was relevant) that would generate ∼71% correct performance for each rat. By targeting 71% correct performance, rats were not guessing, but were well below the performance achieved after learning the IDS (median ± SE: 92.3 ± 2.4%, N = 9 rats in visual modality; 89.3 ± 1.5%, N = 6 rats in auditory modality). Discrimination ability varied across individual rats (**Figure S1**) but the staircase procedure estimated the NoGo stimulus needed to produce a similar degree of discrimination difficulty (71% correct) across rats. In the last 600 trials of the staircase procedure, the percent correct responses were close to the target of 71% (median ± SE: 71.5 ± 1.7% in visual modality and 73.0 ± 1.0% in auditory modality). A Bayesian Wilcoxon test suggested that these data moderately support the null hypothesis that observed performance did not differ from 71% (BF_10_ = 0.210). After obtaining a baseline learning timepoint during an IDS and then requiring either easy or difficult discriminations in the relevant modality for two sessions, in the final step of the experiment we compared the number of trials required to learn an extra-dimensional shift (EDS) against the baseline learning timepoint. In the EDS, novel and easily discriminable stimuli were presented in both modalities and the rats learned to discriminate stimuli in the previously irrelevant modality.

Finally, the three steps of the experiment (i.e., IDS, attention manipulation, EDS) were repeated once and subjecting each rat to the other discrimination difficultly level. This design enabled a within-subjects comparison of EDS learning after easy versus difficult discriminations in the previously relevant modality. One potential confound of this design is that the second EDS could be faster than the first EDS because both modalities had been previously relevant during the second EDS; however, this is unlikely due to the use of novel stimuli in each shift. Moreover, effects of EDS stage were mitigated by counterbalancing (8 rats performed difficult discriminations prior to first EDS and 7 rats performed easy discriminations prior to first EDS). Our data also demonstrate that learning timepoint did not differ between the first and second EDS (median ± SE: 1898 ± 1028 versus 3403 ± 1400 trials, N = 15 rats). A Bayesian within-subjects Wilcoxon test (BF_10_ = 0.557) suggested that the evidence provided weak support for the null hypothesis. Overall, high performance was achieved in the final session of each EDS (median ± SE: 88.9 ± 1.1% in the visual modality and 91.5 ± 0.6% in the auditory modality).

### Difficult discriminations evoked perceptual learning and was associated with increased modality-specific corticocortical neuronal population interactions

We first assessed whether requiring difficult discriminations engaged perceptual attention to a greater extent, relative to easy discriminations, prior to the EDS. **Figure 2A** plots d’ from the last 600 trials of the staircase procedure, through the two attentional manipulation sessions, and in the 200 trials before the introduction of novel stimuli during the EDS. During the staircase procedure, the near-zero effect size demonstrates that the level of stimulus discriminability (easy or difficult) was challenged to similar levels in all rats (**Figure 2B**). However, during subsequent sessions, d’ was higher when the rats were presented with the easily discriminable Go and NoGo stimuli learned during the prior IDS and d’ was lower when the NoGo stimulus was made less distinguishable from the Go stimulus (negative effect sizes, **Figure 2B**).

**Figure 2.**
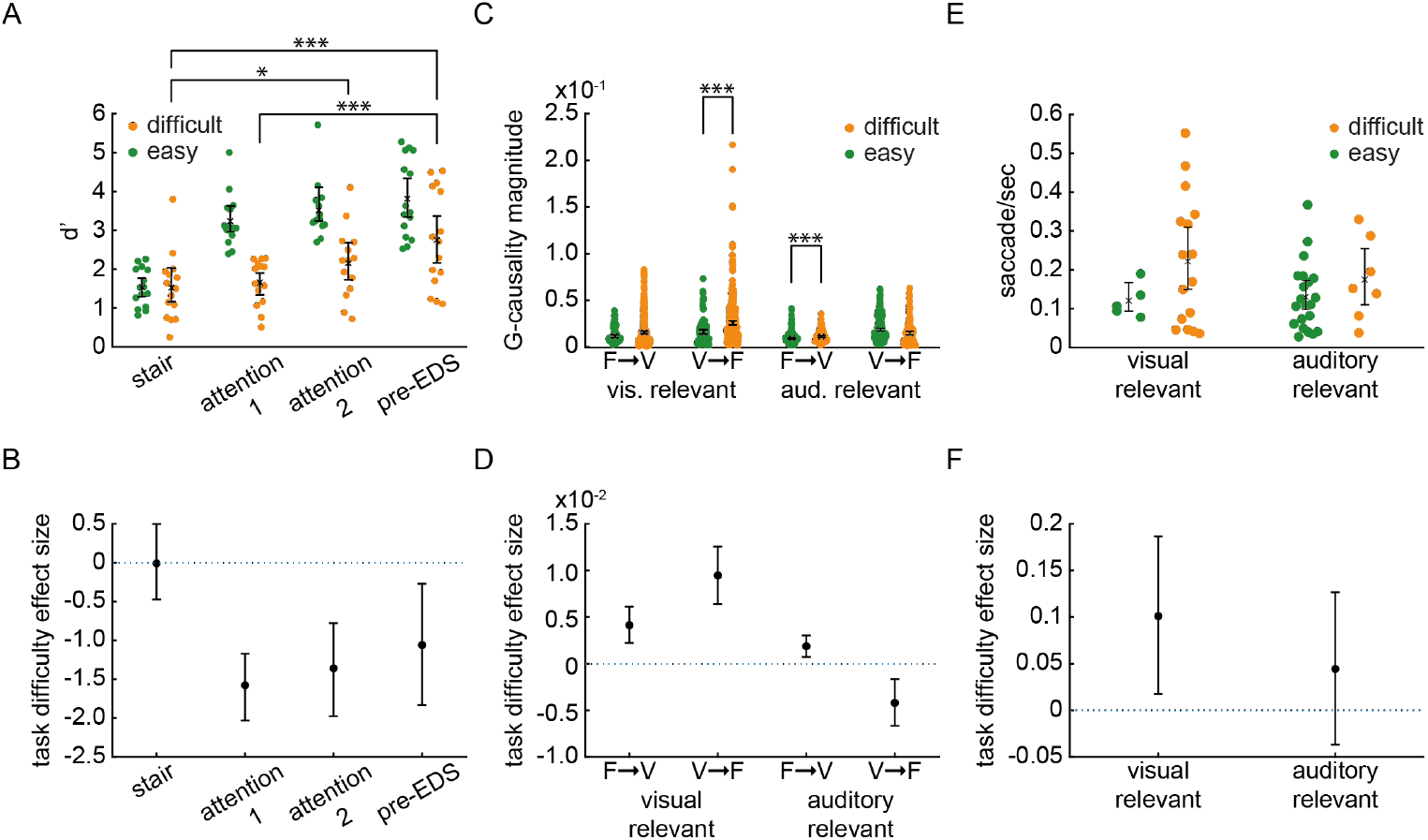
Discrimination ability improved, frontal-sensory cortex EEG Granger causality magnitude increased, and saccade rate increased during repeated sessions of difficult discriminations. **(A)** d’ is plotted in the final 600 trials of the staircase procedure, across two attention manipulation sessions and during the 200 trials before the introduction of novel stimuli in the EDS. The “x” marks the mean performance with standard error in each condition. The dots are rats (N = 15 rats, each rat exposed to both attention conditions; exceptionally, one rat was tested in only one session in the easy condition and another rat was tested in only one session in the difficult condition). Asterisks mark the result of a Bayesian Wilcoxon tests indicating that the alternative hypothesis (i.e., d’ differs between sessions in the difficult discrimination condition) is strongly (*** BF_10_ > 10), moderately (** BF_10_ > 3), or weakly (* BF_10_ > 2) supported by the data. **(B)** Effect sizes between the easy and difficult discrimination conditions (with 95% confidence intervals) are plotted for each stage of the experiment. **(C)** Granger causality magnitude was calculated for all electrode-pairs and collapsed across sessions and rats. The “x” and error bars indicate the mean and standard error in each condition. Each data point is an electrode-pair. The number of electrode-pairs differs due to removal of electrodes with noise contamination (visual relevant – easy: N = 144 electrode-pairs, difficult: N = 432 elec-trode-pairs; auditory relevant – easy: N = 288 electrode-pairs, difficult: N = 144 electrode-pairs. Asterisks mark the result of a Bayesian Wilcoxon tests indicating that the alternative hypothesis (i.e., that magnitude differs between the easy and difficult discrimination conditions) is strongly (*** BF_10_ > 10), moderately (** BF_10_ > 3), or weakly (* BF_10_ > 2) supported by the data. **(D)** The effect sizes (±95% confidence intervals) comparing Granger causality magnitude in the easy and difficult discrimination conditions. **(E)** The saccade rate is plotted for individual rats. These data were not recorded in the entire sample (auditory relevant – easy: N = 21 sessions; visual relevant – easy: N = 5 sessions; auditory relevant – difficult: N = 7 sessions; visual relevant – difficult: N = 16 sessions). **(F)** The effect sizes (±95% confidence intervals) are plotted using same conventions as the other panels in the figure.

We observed a gradual improvement in discriminability across difficult discrimination sessions. A Bayesian two-way ANOVA strongly suggested that performance improved across sessions in the difficult discrimination condition (interaction between session number and discriminability of the stimuli, BF_10_ = 56.802). Post-hoc Bayesian Wilcoxon tests suggested that these data provide weak support of the alternative hypothesis that discriminability improved between the staircase procedure and the second attention manipulation session (BF_10_ = 2.716) but strongly support the alternative hypothesis for an improvement between the staircase procedure and the first 200 trials of the EDS session (BF_10_ = 70.880). There was also strong support for improved discriminability between the first attentional manipulation session and the first 200 trials of the EDS session (BF_10_ = 399.407). Improved discrimination with experience (i.e., perceptual learning) during repeated sessions of difficult discriminations suggests that perceptual attention was more engaged in that condition relative to when easy discriminations were required.

Prior work in non-human primates has shown that, when cued to attend to a stimulus within a receptive field relative to attending outside the receptive field, there is an increase in Granger causality between frontal cortex and visual cortex field potentials recorded in that receptive field r^28^. Thus, increased frontal-visual Granger causality magnitude in indicative of perceptual attention being engaged to improve stimulus discriminability. Therefore, we tested the hypothesis that difficult discriminations were associated with higher frontal-visual cortex Granger causality compared to easy discriminations, but only when the visual modality was relevant. In 4 of the 15 rats, EEG signals were recorded bilaterally across the entire cortex using a flexible 32 electrode array chronically implanted directly onto the skull. Twelve electrodes covered visual cortex bilaterally and four covered frontal cortex bilaterally. Granger causality was measured between all electrode-pairs. We assessed differences at the electrode-pair level; a subject-level analysis was not possible because easy and difficult discrimination conditions were in different modalities for each subject (visual relevant – easy: N = 1 rat, difficult: N = 3 rats; auditory relevant – easy: N = 2 rats, difficult: N = 1 rat). Electrode-pair Granger causality magnitudes were pooled across rats and across the two attention manipulation sessions and the 200 trials prior to the introduction of novel stimuli prior to the EDS.

There were distinct differences in frontal-visual Granger causality magnitude during difficult discriminations relative to easy discriminations depending on which modality was being discriminated (**Figure 2C, 2D**). A Bayesian two-way ANOVA suggested that these data provide strong support for the alternative hypothesis of an interaction between task difficulty and relevant modality (BF_10_ = 8.349E4). Post-hoc Bayesian Wilcoxon tests indicated evidence strongly supporting the alternative hypothesis that bottom-up (visual-to-frontal) Granger causality magnitude was higher during difficult discriminations relative to the easy discriminations when the visual modality was relevant (BF_10_ = 298.155), whereas the null hypothesis was strongly supported in the top-down (frontal-to-visual) direction (BF_10_ = 0.089). On the other hand, when the auditory modality was relevant, the directionality was reversed in that there was strong support for the alternative hypothesis in the top-down direction (albeit, weaker that when the visual modality was relevant, BF_10_ = 16.287) while the null hypothesis was moderately supported in the bottom-up direction (BF_10_ = 0.037). These data suggest that requiring difficult discriminations in one modality alters interactions between frontal cortex and the sensory cortex associated with the modality being discriminated.

Finally, we used saccade rate as a behavioral measure of attentive behavior. In the difficult discrimination condition, rats performed more saccades per second especially when the visual modality was relevant (**Figure 2E, 2F**). There was moderate support for the alternative hypothesis that difficult discriminations were associated with increased saccade rate (BF_10_ = 3.084).

### Engaging perceptual attention onto one modality slows extra-dimensional set-shifting

We predicted that engaging perceptual attention onto one modality would affect future outcome-based learning. Specifically, we tested the hypothesis that difficult discriminations would increase the shift cost (i.e., make learning responses to novel stimuli using the previously unrewarded modality during an EDS more difficult relative to baseline learning of responses to novel stimuli using the previously rewarded modality during an IDS). We found that the prior experience performing difficult discriminations in one modality was associated with an increased shift cost for learning about novel stimuli in the other modality (**Figure 3**). The effect size was large: on average 1,726 more trials were required for learning about novel stimuli after performing difficult discriminations in the other modality. The 95% confidence intervals of this effect were a minimum of 577 additional trials required for learning and up to 2,874 additional trials. A Bayesian within-subjects Wilcoxon test indicated strong support for the alternative hypothesis (BF_10_ = 24.345). This finding was robust against changes in the bin size using for calculating % correct performance and the definition of the learning timepoint. A Bayesian within-subjects Wilcoxon test indicated the data provide strong support for the alternative hypothesis when 10, 20, 30, and 100 trial bin sizes were used with the 80% correct learning timepoint (BF_10_ = 20.145, 25.070, 11.348, and 21.207, respectively). Similarly, when considering instead 75% correct performance as the learning timepoint, data across all trial bin sizes (10, 20, 30, 50, and 100 trials) strongly supported the alternative hypothesis (BF_10_ = 16.737, 25.039, 13.790, 18.155, and 22.633, respectively). Although there was a clear effect of discrimination difficulty in the previously relevant modality on subsequent learning of the EDS, the degree to which perceptual learning occurred in the previously relevant modality (change in d’) was not correlated with the shift cost (Pearson’s R = -0.334; Bayesian Correlation BF_10_ = 0.169 which indicated moderate-to-strong support for the null hypothesis). Our data suggest that difficult discriminations in one modality subsequently slows learning the responses associated with novel and easily discriminable stimuli in another modality.

**Figure 3.**
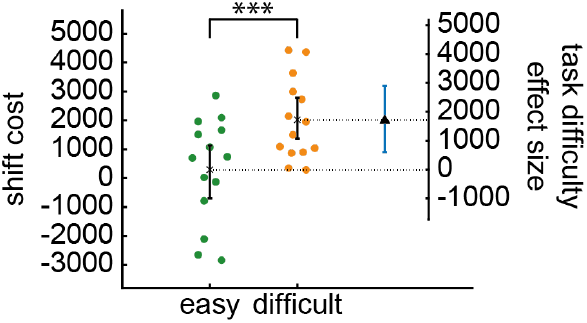
Slower EDS learning after difficult discriminations in the previously relevant modality. The plot shows the shift cost when the EDS occurs after requiring easy or difficult discriminations in the previously relevant modality. The dots are individual rats (same rats in both conditions). The inset shows the effect size (±95% confidence intervals). One rat was removed because it was an outlier in the difficult discrimination condition (2.6 standard deviations from the median) since it performed in an opposing trend to the other rats. However, including the outlier, the alternative hypothesis was still weakly supported, whereas the null hypothesis was not (BF_10_ = 2.038). Asterisks mark the result of a Bayesian Wilcoxon test indicating that the alternative hypothesis is strongly (*** BF_10_ > 10), moderately (** BF_10_ > 3), or weakly (* BF_10_ > 2) supported by the data.

### Number of rewards, arousal, and response criterion were not affected during the manipulation of perceptual attention

We propose that learning was affected by the historical focus of perceptual attention, but other non-attentional factors may differ between task conditions requiring easy or difficult discriminations and those other factors could influence subsequent learning of the EDS. For instance, rats could obtain fewer rewards during difficult discriminations by committing more response omissions. However, we found that a similar number of rewards were obtained in both discrimination conditions (**Figure 4A**). A Bayesian Wilcoxon test indicated either a lack of evidence supporting either the null or alternative hypotheses during the first attentional manipulation session (BF_10_ = 1.939), weak support for the null hypothesis during the second attentional manipulation session (BF_10_ = 0.574) and modulate support for the null hypothesis during the 200 trials prior to introduction of novel stimuli during the EDS (BF_10_ = 0.385). Moreover, the effect sizes were small (**Figure 4B**). Rats received only on average 7.2% fewer, 9.5% fewer, and 3.2% fewer rewards during difficult discriminations in those task epochs.

**Figure 4.**
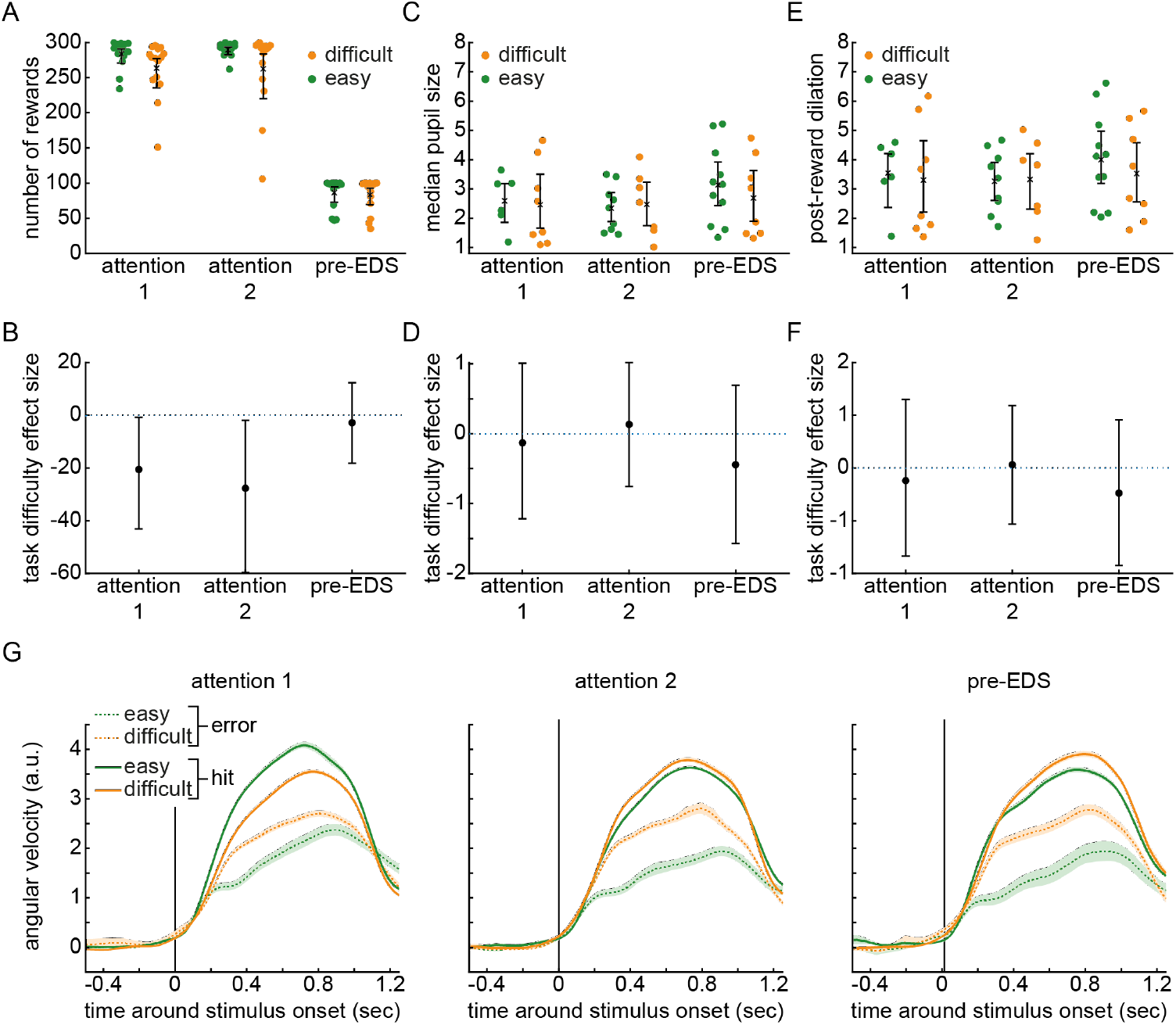
Non-attentional factors did not differ between easy and difficult discrimination conditions. **(A)** The number of rewards obtained in different stages of the task. Each dot is a rat (N = 15 rats tested in both conditions). **(B)** The effect sizes (with 95% confidence intervals) compare the number of rewards received in the difficult discrimination versus easy discrimination condition at each task stage. Differences are small magnitude. **(C)** The median pupil size is plotted for each rat and each session, comparing the difficult discrimination and easy discrimination conditions. The units are arbitrary and scaled by 10^−3^. Pupil size was measured in a subset of rats. Each dot indicates the session median pupil size for one rat. **(D)** The effect sizes (with 95% confidence intervals) compare the difference in median pupil size between easy and difficult discrimination conditions at each task stage. **(E, F)** The post-reward pupil dilation magnitude is plotted. Plotting conventions are identical to those in C and D. **(G)** The plots show treadmill velocity traces aligned to stimulus onset in the different task stages. The units are rotary encoder voltage change/sec scaled by 10^3^. The lines are the mean across rats and the shading is the standard error of the mean. Solid lines represent velocity on hit trials. Dotted lines represent error (false alarm) trial running. Most groups contain 11 rats except for 10 rats in the difficult discrimination condition for the second attention manipulation session and 12 rats in the difficult discrimination condition for the 200 trials preceding the introduction of novel stimuli before the EDS.

Another factor that could be altered during the manipulation of perceptual attention is arousal. We used pupil size as an index of arousal (**Figure S2A**). Luminance was held constant across subjects and task sessions by placing the experimental set-up inside of a walk-in, sealed faraday cage and using a constant head-fixation location. Rats were transported in light-blocking cages and tested under red (>590 nm) light. The median pupil size did not differ between conditions (**Figure 4C, 4D**). A Bayesian Wilcoxon test indicated weak support for the null hypothesis (first attention manipulation session, BF_10_ = 0.470; second attention manipulation session, BF_10_ = 0.439; during the 200 trials prior to introduction of new stimuli during the EDS, BF_10_ = 0.505). The effect sizes were small and in opposing directions across task epochs (means: 5.0% smaller, 5.7% larger, and 14.2% smaller) indicating that arousal was neither consistently higher nor lower when difficult discriminations were required. We also measured arousal related to reward consumption using the peak of the trial-averaged pupil dilation aligned to reward delivery (**Figure S2B**). The reward associated pupil dilation also did not differ between discrimination conditions (**Figure 4E, 4F**). A Bayesian Wilcoxon test indicated weak support for the null hypothesis (first attention manipulation session, BF_10_ = 0.465; second attention manipulation session, BF_10_ = 0.447; during the 200 trials prior to introduction of new stimuli during the EDS session, BF_10_ = 0.462). The effect sizes were small and not in a consistent direction across task epochs (means: 6.8% smaller, 1.9% larger, and 11.9% smaller across task epochs). Overall, these data suggest that arousal did not differ between sessions requiring easy discriminations compared to difficult discriminations before the EDS.

Finally, it is possible that greater error commissions during difficult discriminations was associated with a shift in response criterion and this could influence subsequent learning during the EDS. We predicted that during difficult discriminations, a shifted response criterion would be indicated by a reduced Go response latency (running onset) and an increased Go response magnitude (peak velocity) and that these changes would be in the same direction on hit and false alarm trials. We examined these two aspects of the response by directly inspecting Go response movement dynamics (**Figure 4G**). There were no apparent differences in the latency to initiate responses. In all 3 sessions, during difficult discriminations, false alarm responses were of a higher magnitude. On the other hand, hit responses were lower magnitude during the first attention manipulation session, whereas in the second attention manipulation session and the 200 trials prior to the introduction of novel stimuli before the EDS, the hit responses were slightly higher magnitude. This increased magnitude was not as large as the increase on false alarm trials. In sum, changes in response dynamics during difficult discriminations were inconsistent across task epochs and trial types.

## Discussion

Studies on the role of attention in learning have focused on how manipulating associations between stimuli and rewards can cause attention to shift between stimuli dimensions during learning ^1,2^. However, attention has also been conceptualized as a filter that improves sensory perception ^9–11^. It is unknown whether focusing perceptual attention onto one stimulus dimension could subsequently influence how outcome-based attention is controlled during learning to shift dimensions in the context of novel stimuli. Although engaging perceptual attention with a perceptual learning paradigm has been shown to slow switching from one stimulus dimension to another ^29^, such task switching does not assess how perceptual attention modulates outcome-based learning since it required switching between two known rules using familiar stimuli. Thus, it remains unknown whether engaging perceptual attention interacts with outcome-based attention during learning.

Here, we demonstrate that perceptual attention directly affects learning rate and therefore uncover a new role of attention in learning.

We manipulated modality-specific perceptual attention by requiring either easy or difficult discriminations in the relevant modality prior to testing the ability to learn an EDS to the other modality with novel stimuli in both modalities. We show that, during repeated sessions of difficult discriminations, perceptual learning occurred. This was accompanied by altered modality-specific frontal-visual EEG Granger causality magnitude during difficult discriminations. This result is consistent with prior findings in non-human primates showing that frontal-visual Granger causality increases during the engagement of perceptual attention ^28^. It is also consistent with prior work showing that perceptual learning may engage higher level cortex in the usage of sensory information to control behavior ^30^. We interpret the occurrence of perceptual learning and the accompanied modality-specific cortico-cortical Granger causality increase as indirect signs that modality-specific perceptual attention was engaged during difficult discriminations. There has been some debate over whether observing perceptual learning indicates such engagement ^31^. For instance, perceptual learning occurs in conditions that do not engage top-down perceptual attention, such as presentation of subliminal stimuli or instructing subjects to not attend to stimuli ^32,33^. On the other hand, numerous studies have suggested that perceptual learning requires the engagement of top-down perceptual attention ^17–22^. It is likely that in goal-directed tasks, such as the one used here, perceptual learning is due to attentive practice discriminating the stimuli ^34^. Furthermore, we used saccade rate as a potential measure of attentional engagement. The frequency of this behavior increased during difficult discriminations, which provides additional support for the engagement of attention during difficult discriminations. Overall, our behavioral and electrophysiological results suggest greater engagement of perceptual attention during difficult relative to easy discriminations.

After differentially engaging perceptual attention with either easy or difficult discriminations for two sessions, we presented novel and easily discriminable stimuli in both modalities and assessed learning responses to stimuli in the modality that was previously irrelevant (i.e., an EDS). Importantly, the use of easily discriminable stimuli during the EDS confined manipulation of perceptual attention to pre-learning time points, rather than during the learning process itself. Thus, the historical focus of perceptual attention was manipulated. We found that engaging perceptual attention onto one modality slowed learning about novel stimuli in the other modality relative to a baseline number of trials needed to learn discriminating novel stimuli without changing the relevant modality (i.e., an IDS). Targeting more difficult discriminations or increasing the number of sessions to drive additional perceptual learning could increase the effect on shift cost.

Increased shift cost is evidence for the mental formation of an ‘attentional set’, a rule that classifies complex stimuli according to a single feature ^5,35^. Engaging perceptual attention by requiring difficult discriminations may have increased the formation of an attentional set. In prior work, attentional set formation has been evoked by maintaining the same rewarded stimulus dimension across repeated novel discriminations or by repeating reversals of stimulus-outcome relationships ^5,7,35,36^. These manipulations overtrain the rewarded stimulus dimension. Notably, the two attention manipulation sessions requiring easy or difficult discriminations had an identical number of trials so that one condition did not involve overtraining relative to the other. Thus, in contrast to prior work, our result suggests a new method to generate an attentional set that does not require overtraining stimulus-outcome associations.

It is likely that perceptual attention is one of many cognitive processes that affect attentional set-shifting task performance. For instance, outcome-related attention has a clear role in the task ^1,2^. Given that multiple factors can influence task performance, it is not surprising that the amount of perceptual learning did not correlate with the subsequent shift cost at the individual subject level. However, we were able to exclude several non-attentional factors that could differ between task conditions requiring either easy or difficult discriminations and, thus, affect subsequent learning of the EDS. One such factor, arousal, was assessed using pupillometry and was similar in both discrimination conditions. Reinforcement is another non-attentional factor driving perceptual learning ^37^ but we found that reward consumption did not differ across the task conditions. Finally, we assessed whether response criterion differed by directly measuring the dynamics of the stimulus-guided behavioral response. Although difficult discriminations were associated with changes in response magnitude, these were inconsistent across task epochs and trial types. Additionally, latency changes were not observed. Collectively, the analysis of response dynamics suggests that criterion changes were not a major factor modulating subsequent EDS learning.

Our data suggest that there are two *interacting* types of attention that affect rule learning: first, “how much” attention has been *historically* focused on perceptual features of stimuli ^17^, and second, attentional capture by decision outcomes. The involvement of two types of attention in rule learning may help explain why psychiatric disorders share impaired attentional set-shifting as a trans-diagnostic symptom. For instance, a diagnosis of autism is associated with deficits in EDS learning ^12^. Conspicuously, these individuals also present deficits in perceptual learning and perceptual attention ^27^. EDS learning is also impaired for individuals diagnosed with schizophrenia ^13^, but without the deficits in perceptual learning ^38,39^. Schizophrenia is, however, associated with impaired reinforcement learning because of insensitivity to receipt of rewards ^40,41^. These findings from the psychiatric literature, considered in the context of our findings, suggest a potential explanation for why impaired attentional set-shifting is a trans-diagnostic symptom of different psychiatric disorders. We propose that individuals with schizophrenia may have impaired outcome-based attention, which leads to impaired EDS learning. In contrast, individuals diagnosed with autism may have impaired perceptual attention, providing a different route to impaired EDS learning. If outcome-based and perceptual attention are predominantly weighted toward separate neural circuits, then the treatment of impaired attentional set-shifting may require targeting different neural circuits in autism versus schizophrenia.

Understanding the neurobiology of behavior requires first defining which cognitive functions are in use during a behavioral task ^42,43^. During the attentional set-shifting task, outcome-related attention constrains how values are calculated and updated ^1^. Based on our finding that focusing perceptual attention onto one modality impairs subsequent learning in another modality, we propose that, in addition to outcome-related attention, the historical focus of perceptual attention places an additional constraint on learning in the attentional set-shifting task. Prior work has shown that engaging perceptual attention has an associated cost, in that it can cause subjects to imagine stimuli that are not physically present ^44^. Here, we show yet another cost of perceptual attention, which is on subsequent learning about novel stimuli. Although it may be beneficial for survival when attention sharpens perception of one sensory modality, in the context of learning this may occur at the expense of another modality and contribute to inflexible behavior. This double-edged sword of attention illustrates a cognitive constraint placed on learning.

## Acknowledgements

We thank Prof. Lewis Chuang and Prof. Joshua Gold for discussions on psychophysics and perceptual learning. We appreciate guidance from Prof. Laura Busse and Dr. Steffen Katzner on pupillometry measurements in rodents. We are grateful for comments on the manuscript from Prof. Peter Dayan, Prof. Luigi Acerbi, Prof. Anna Levina, Prof. Marie Carlén, and Dr. Aron Koszeghy. We thank Prof. Geoffrey Schoenbaum and Dr. Marios Panayi for insightful discussions regarding learning theory. We wish to thank the Finnish Grid and Cloud Infrastructure (FGCI) for supporting this project with computational and data storage resources. This work was funded by the Max Planck Society and the University of Helsinki (Helsinki Institute of Life Science). This paper was typeset with the bioRxiv word template by @Chrelli:www.github.com/chrelli/bioRxiv-word-template

## Author contributions

Conceptualization – NT; Formal analysis – DV, NT; Funding acquisition – NKL, NT; Investigation – DV, NS, RI, HB, ISZ, Methodology – DV, RI, MW, NT; Project administration – NT; Resources – NKL; Supervision – NT; Visualization – DV, NT; Writing – DV, NT.

## Materials and Methods

### Subjects

Male rats (Lister-Hooded) were obtained from Charles River at a weight of 140 grams to 190 grams. Rats were housed in pairs for a 7-day acclimation period prior to implantation with a chamber and head-post and, in some cases, an EEG array. After implantation, rats were single housed. All behavioral testing was carried out during the rats’ active phase and housing illumination was between the hours of 7PM and 7AM. All procedures were carried out with the prior approval of local authorities and in compliance with the European Community Guidelines for the Care and Use of Laboratory Animals.

### Surgical procedures

The rat was anesthetized using isoflurane (induction chamber for 4% for 3 min and 2.5% for 5 min followed by 2.5% or less via the nose cone). Anesthetic concentration was adjusted throughout the procedure to maintain a heart rate of ∼300 to 350 beats per minute. The rat was head-fixed using ear bars. After fixation, the rat was with buprenorphine (0.06 mg/kg, s.c.), meloxicam (2.0 mg/kg, s.c.), and enrofloxacin (10.0 mg/kg). We injected lidocaine (0.5%, s.c.) under the scalp. Surgical procedures began as soon as the rat was not responsive to the paw pinch (usually 10 to 15 minutes after the injection of painkillers). Skin and underlying connective tissue were removed to expose most of the frontal bone and laterally and posteriorly to the surrounding musculature. Soft tissue bleeding was stopped using cauterization. Skin and underlying connective tissue were removed to expose most of the skull from the frontal bone continuing posteriorly to the neck muscle and also laterally to the left and right temporal muscles. The bone was wiped dry and cleaned with 5% H2O2 applied using a cotton swap. The H2O2 was immediately removed by washing with 40 mL of saline. At this stage, for a sub-set of rats, an EEG array (Neuronexus, CM32) was laid onto the skull and fixed in place using dental cement (2-stage, powder/liquid Paladur). The surrounding exposed bone was scratched using bone curette in a grid pattern to increase the adhesion of the subsequently applied UV-curing primer that serves as a base for fixation of the chamber and head-post onto the skull. If no EEG array was implanted, then the entire skull was scratched. Primer was applied using a brush (Omnibrush, DentalBauer) and UV cured for 30 sec at full intensity (Superlite 1300, M+W Dental). A custom-made skull implant was used for head-fixation (machine shop, Max Planck Institute for Biological Cybernetics). The implant was fixed onto the skull using UV-curing dental cement (Tetric Evoflow, Dental Bauer) that bonds to the underlying layer of primer on the skull. A craniotomy was made on the left occipital bone for a ground wire. The ground wire was 99.9% pure silver. One end was flattened using an industrial press into a ∼1-2 mm wide rectangle, which then was twisted into a roll to fit the craniotomy and inserted into the space between bone and dura. A rolled shape was used to increase the potential surface area in contact with CSF. The craniotomy was filled with viscous agar, which stabilized the wire and is also conductive. The other end of the ground wire was soldered to the ground wire of the electrode interface board of the EEG array. The wires and array were buried under the thick layer of dental cement (Paladur). The skin around the implant was glued to the implant using tissue glue (Histoacryl, B. Braun). Post surgical recovery lasted five days. During the first three days (surgery itself was counted as day one), the animal was injected either every 12 hours with buprenorphine or every 24 hours with meloxicam (same dosages as pre-operative). During the first five days, enrofloxacin was injected every 24 hours (same dosage as pre-operative). A rehydrating, nutritious, easily consumed, and palatable food was provided (DietGel Recovery, Clear H2O).

### Handling and habituation

Rats were handled daily for at least five minutes per day from the day of arrival in housing until the day of surgery, which was 7 days. After surgery, the rats were water restricted for at least one full day prior to the first head fixation. Water restriction procedures are explained in detail in the next section. The training procedure can be divided into habituation, training of instrumental response (head-fixed treadmill running), training of stopping response (suppression of premature responding), and training of a NoGo response and discrimination of sensory stimuli using a Go/NoGo response paradigm.

Habituation consisted of one day, which included head fixation on the treadmill with a reward port aligned to rat’s mouth and visual confirmation that it could be easily licked. Extremely tiny movements on the treadmill, such as actions that maintained balance, were rewarded with a 5 uL drop of 10% sucrose solution. The first session lasted 20 min regardless of the amount of water consumed.

Starting from the second day, we trained the instrumental response. The next 1 to 3 sessions were identical to the habituation day, except that every reward was accompanied with a 0.1 sec duration tone (500 Hz). The tone served as a bridging stimulus to help the rats learn the association between movements and rewards. Each reward delivery was 5 microliters. We gradually increased the velocity threshold to encourage stronger movements, first consisting of large swinging motions of the entire body and eventually stepping. Each session lasted 30 min or until they consumed 8 mL of sucrose water, whichever occurred first. When the rat was running and licking simultaneously over continuous periods lasting ∼5 to 10 seconds, they typically received 7 mL of sucrose water during the behavioral session. At this stage, the training protocol was advanced.

In the next training stage, the Go stimulus was introduced (either auditory or visual, see results section for details). Initially, the stimulus duration was 15 sec with an inter-trial interval (ITI) (selected randomly from a 2 to 3 sec distribution). Rats were trained to run during the stimulus. Any threshold crossing movement that occurred during the stimulus was rewarded and repeated threshold crossings during the same stimulus presentation provided repeated rewards. Threshold crossing movements were defined by a distance threshold which was the same for all rats and corresponded to a burst of running (∼4 strides). Normally, rats required one session (∼30 minutes long) to both run during the stimulus and avoid running during the ITI. This indicated that the rat had learned to associate the stimulus with the previously trained action-outcome association.

The next phases of training focused on training the stopping response. We first introduced a timeout (0.5 sec duration) after a premature response, which was defined as running over a velocity threshold during the ITI. Premature responses also resulted in restarting the ITI after the timeout ended. The velocity threshold was identical for all rats and corresponded to a single step on a treadmill. The premature timeout and restarting of the ITI remained in the task design throughout all subsequent stages of the experiment. After one to two sessions, the rats learned to suppress premature running. At this point, stimulus duration was reduced to 5 seconds and the ITI was reduced to 1 to 2 seconds. Typically, after five sessions, rats were responding to nearly every stimulus and had adjusted their responding to soon after the onset of the stimulus due to the limited window to obtain reward. At this stage, the stimulus duration was further reduced to 2.5 seconds and the ITI was reduced to 1 to 1.5 seconds. Reward drop size was increased to ∼8 uL. We trained rats for 2 or 3 sessions at this stage to adapt them to the shorter stimulus, but there was no qualitative change to their behavior between the 5 sec and 2.5 sec stimulus. After 2 or 3 sessions, the stimulus duration was reduced to our target duration of 1.5 seconds and the ITI was changed to 0.5 to 1.0 sec. Reward drop size was increased to 10 uL. Given the brief stimulus duration, the task became a speeded response task. Stable behavior at this stage was defined as an omission rate of less than 10% and water consumption of at least 6 mL per session.

As soon as behavior was stable at this stage, the Go/NoGo paradigm was trained. A NoGo stimulus was introduced. Due to the pre-potent drive to respond, we introduced a response offset window (0.75 sec after stimulus onset), during which running did not count toward the distance threshold. This allowed low latency movements, but forced the rat to perceptually appraise the stimulus and then make a decision. Responding to the NoGo stimulus (i.e., crossing the distance threshold) resulted in a timeout of 6 seconds. Upon response threshold crossing, the stimulus was extinguished and the timeout began. A correct response to the Go stimulus also resulted in extinguishment of the Go stimulus. Reward was delivered in the form of three drops (10 uL) each separated by 0.5 sec. While the duration of prior training sessions previously was based on subjective judgement about the satiety of the rat, the Go/NoGo paradigm training always lasted 400 trials. Performance (number of hits and correct rejections out of total number of trials) reached 80% after 2 to 6 sessions. After reaching this criterion levels of performance, the irrelevant stimulus modality was introduced. Training was continued with the compound auditory-visual stimuli until performance was >85%. Typically, this required between 2 and 10 sessions of training.

An equal number of Go and NoGo stimuli were presented in each session. The two irrelevant modality stimuli were each presented an equal number of times with the Go and NoGo stimuli. We also introduced a requirement that no more than three NoGo trials could occur in succession due to difficulty suppressing responses. Three NoGo responses would require sitting immobile for a 0.5 to 1.0 sec ITI, then a 1.5 sec NoGo stimulus and repeating this two more times. Although it was possible for rats to do this, we determined that it was an unreasonable level of difficulty.

Upon reaching this criterion levels of performance, training was completed and rats started the experiment consisting of two baseline sessions followed by an intra-dimensional shift. Details of the experimental paradigm are provided in the results section.

### Water restriction

During the handling procedure, the availability of water was restricted. During training, rats were provided 8 to 12 mL of water per day (including the volume consumed during the behavioral task) for 5 days followed by 48 hours of ad libitum water availability. We initially provided 10 mL of daily water. In rare cases where body weight loss was approaching 15% drop from the last weight during ad libitum water availability, then 12 mL of daily water was provided. The omission rate of some rats was high unless only 8 mL of water was provided per day. After training in the basic Go/NoGo task, rats were switched to a continuous schedule of water restriction, without intermittent 48-hour windows of ad libitum water availability to prevent fluctuations in motivation caused by intermittent changes in water availability, as well as a stress response due to intermittent water availability (Vasilev et al., 2021), which could affect task performance.

### Attentional set-shifting task

After training in the Go/NoGo task, rats performed a series of intradimensional and extra-dimensional set-shifts in which completely novel stimuli were presented and the rats needed to learn the new discrimination through trial-and-error. All rats performed an IDS followed by an EDS and then the procedure was repeated, each time with novel stimuli.

Each IDS was preceded by two baseline sessions to ensure stability of learning the prior discrimination. After learning an IDS, it was followed by a psychophysics experiment that aimed to determine the attribute of the NoGo stimulus that would generate 71% correct performance (see next section for description of the psychophysics experiment).

Each EDS was preceded by two attention manipulation sessions. If the manipulation was to maintain a normal level of attention, then these two sessions were identical to baseline sessions, in that rats needed to demonstrate their ability to perform the previously learned discrimination from the IDS. If the manipulation aimed to hyper-focus attention, then during these two sessions the rats were presented with the Go stimulus from the previously learned IDS and the individually tailored NoGo stimulus designed to generate 71% correct discrimination performance. In both cases, the same stimuli from the irrelevant modality, which were presented during the IDS, were presented during attentional manipulation.

During the IDS or EDS, the new stimuli were introduced at trial number 200, such that the first 100 trials of the session served to confirm performance of the previously learned rule. Rats were allowed as many behavioral sessions as needed to finish learning. Each session lasted for 600 trials. Completed learning was defined >85% correct performance for one session.

### Sensory stimuli

Stimuli in the visual modality were black and white drifting gratings (spatial frequency = 0.005 cycles/pixel). The visual Go and NoGo stimuli were 70 deg apart. Visual stimuli were presented 16cm from the rat’s head. Auditory stimuli ranged from 0.5 kHz to 64.339 kHz (sampling rate = 192 kHz) and the Go and NoGo stimuli were three-quarters of an octave apart. Auditory stimuli were presented using a Lynx E44 sound card (192 kHz sampling rate), an amplifier with a 0.1 kHz to 100 kHz range (Yamaha, AX-397), and stereo speakers with a 0.004 kHz to 100 kHz range (Sony, MDR-Z7) mounted on either side of the rat’s head. A microphone (Bruel & Kjaer, 4959-A) with a range of 0.004 kHz to 70 kHz was used to confirm sound delivery and tune stimulus amplitudes so that the sounds were all ∼65 dB.

### Staircase procedure

The psychophysics task used a 2-down / 1-up procedure, whereby each correct response resulted in the NoGo stimulus changing toward the Go stimulus by 2 steps and each false alarm resulted in moving the NoGo stimulus 1 step away from the Go stimulus. The step size for auditory stimuli was defined as 100 steps in logarithmic space between the Go and NoGo stimulus and for visual stimuli the step size was 0.5 degrees. Rats were run on the psychophysics task until they could no longer improve their performance for most of a behavioral session (600 trials). This typically required 2-3 sessions.

### Pupillometry data acquisition and processing

Videos were recorded at 45 frames per second from the rat’s right eye with near-infrared illumination (Thor Labs LED, M850L3 and Thor Labs Collimation optics, COP4-B). Frames were acquired using a near-infrared camera (Allied Vision, G-046B) and variable zoom lens, fixed 3.3x zoom lens, and 0.25x zoom lens attachment (Polytec, 1-60135, 1-62831, 6044). Acquisition occurred over a GigE connection (MATLAB image processing toolbox). The camera provided a TTL pulse with each video frame. These TTL pulses were recorded directly into the neurophysiology and behavior control system (Neuralynx).

We used an in-house custom algorithm and computer code, implemented in Python, to extract pupil size from the recorded video frames. Before processing the sequence of frames, one of them was randomly selected to manually crop the rat’s eye to specify the region of interest for the algorithm. At this stage, several meta parameters for each of the following steps were simultaneously selected using manual inspection in a GUI (**Figure S3**). This was left intentionally to an operator’s judgement due to natural noise in the image of the rat’s eye (e.g., the eyelid, lashes, or fur could cast a shadow on the pupil). First, each cropped image was blurred using Gaussian blur. The image was then converted into a binary black and white image. The edges on this image were defined using the Canny’s algorithm (Canny, 1986). Then, closed contours were selected using an algorithm described by Suzuki and colleagues (Suzuki and Abe, 1985). Finally, ellipses were fitted into closed contours using the method of Fitzgibbon and colleagues (Fitzgibbon and Fisher, 1995). If there happened to be several elliptical contours, the ellipse with minimal eccentricity was selected. In cases where the algorithm was not able to find an ellipse of an area bigger than predefined minimal allowed area (or smaller than maximal, respectively), then the value of the pupil in this frame was left blank. This was also applied to the frames which captured the animal blinking. Blank frames were linearly interpolated. The pupil detection algorithm was implemented using the OpenCV package in Python 3.7.

### EEG signal acquisition

EEG signals were recorded using a flexible polyimide array with 32 platinum electrodes (Neuronexus, H32). Signals were recorded against animal ground, pre-amplified at the rat’s head (Neuralynx, HS-36), and then amplified and digitized at 32 kHz (Neuralynx, Digital Lynx SX). The analysis focused on bilateral electrodes placed over frontal cortex (locations relative to Bregma: 1.5 mm anterior, ±1.2 mm lateral; 3.6 mm anterior, ±1.2 mm lateral) and visual cortex (locations relative to Bregma: 5.0 mm posterior, ±1.5 mm lateral; 5.0 mm posterior, ±3.0 mm lateral; 5.0 mm posterior, ±4.4 mm lateral; 7.0 mm posterior, ±1.5 mm lateral; 7.0 mm posterior, ±3.0 mm lateral; 7.0 mm posterior, ±4.4 mm lateral).

### Treadmill velocity signal acquisition

The treadmill was a freely rotating fiberglass cylinder that could rotate freely either forward or backward on a metal axis inserted into low friction ball bearings. Treadmill velocity was calculated from an analog signal corresponding to the rotation of the axis, which was recorded using a rotary encoder (US Digital, MA3-A10-125-B). The analog voltage output of the rotary encoder varied between 0V and +5V mapped onto 0 to 360 degrees. The signal was recorded as an analog input in the Neuralynx amplifier. The rotational angle signal was sampled at 32 kHz. It was unwrapped in MATLAB to grow continuously by differentiating the signal and detecting the 0V-to-5V (0-to-360 degrees) and the 5V-to-0V (360-to-0 degrees) resets using the MATLAB function, peakdetect. After unwrapping the rotational angle over time, the signal was lowpass filtered (5 Hz), then downsampled to 100 Hz, and finally differentiated to obtain the angular velocity.

### Data analysis

We defined the shift cost as the number of trials required to reach the learning point in the EDS minus the number of trials required to reach the learning point in the preceding IDS. The learning point was defined as the trial when the performance reached 80%. We defined performance by fitting a “performance curve” to the proportion of correct responses in windows of 50 trials calculated by sliding the window in steps of one trial. The performance curve was obtained by fitting a sigmoid function to the proportion of correct responses using the psignifit package with the sigmoid option set to cumulative Gaussian and experiment type set to 2AFC (see https://github.com/wichmann-lab/psignifit/wiki for details).

Granger causality was calculated using the MVGC toolbox for MATLAB (Barnett and Seth, 2014). Standard parameters were used. Before calculating Granger causality, the EEG signals were downsampled to 100 Hz. Calculations were made on the signal from the entire behavioral session without respect to the trial structure of the task. Each signal-pair was conditioned upon a third electrode, moving through all 16 other electrodes on the array that were not located over frontal or visual cortex. We reported the Granger causality magnitude for a frontal-visual electrode-pair as the average magnitude across all 16 conditioning channels.

### Statistics

We used estimation statistics and report effect sizes and the confidence intervals for effect sizes (DABEST toolbox in MATLAB (Calin-Jageman and Cumming, 2019; Ho et al., 2019)). Bayesian statistics were used for assessing evidence (or lack thereof) for the null hypothesis and for the alternative hypothesis (Keysers et al., 2020). Bayesian statistics were calculated in JASP software. The choice of Bayesian test was selected based on whether the data were normally distributed, which was assessed using a Kolmogorov-Smirnov goodness-of-fit hypothesis test (kstest, MATLAB).

---

**Figure S1:**
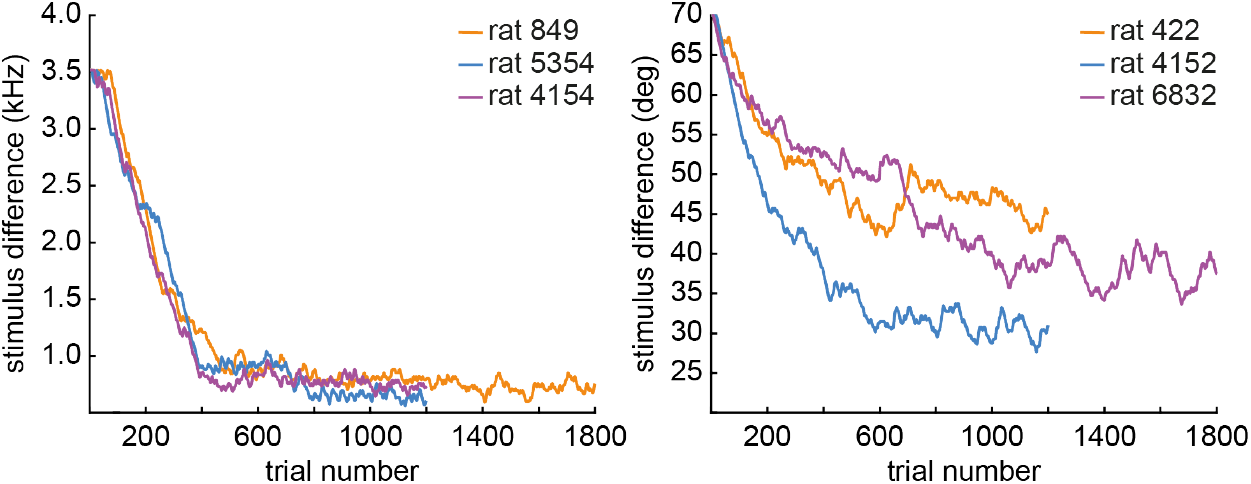
The parameter difference between Go and NoGo stimuli during the staircase procedure. Data are shown for 6 example rats in the auditory (kHz) or visual (deg) modality.

**Figure S2:**
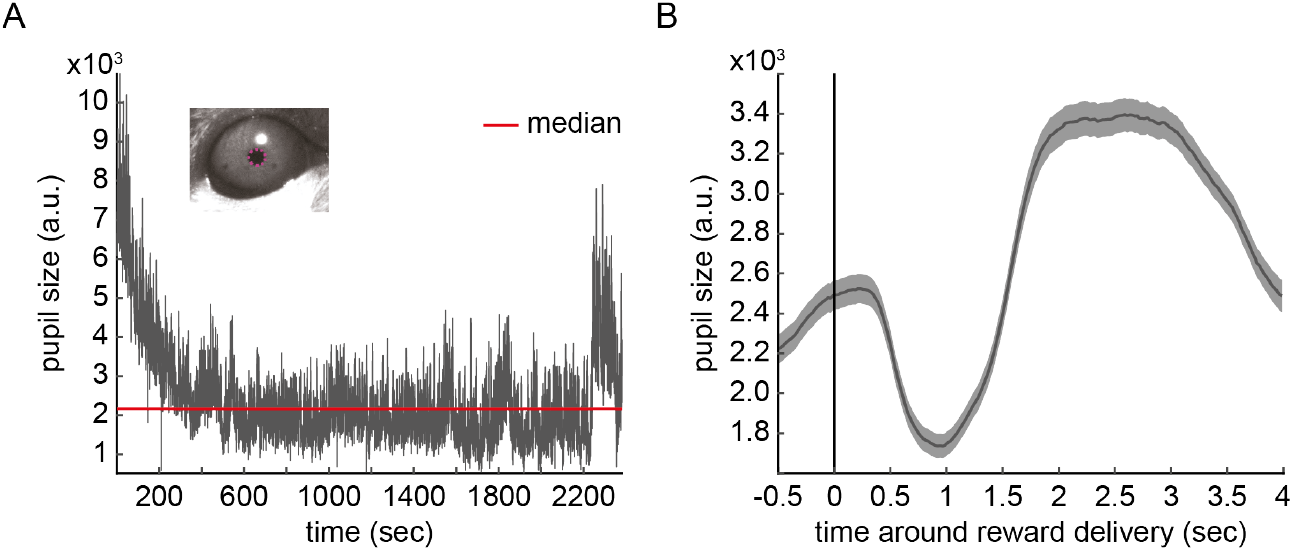
The pupil size is shown across an entire example session and in a brief time window aligned to reward delivery. **(A)** Pupil size is plotted over the example session from a single rat. Pupil size was larger at the start of the task across sessions and rats, even when a task is not performed and in stable conditions of darkness. The median is marked by a red line. The inset shows a single frame of the video with the pupil size indicated by the magenta elipse. Pupil size was extracted from video of the eye recorded at 45 frames per second. **(B)** The trial-averaged peri-reward pupil size aligned to reward delivery is plotted for one example session. The shading shows the error across trials. Post-reward dilation was the peak of the trial averaged pupil dilation during the 4 sec window after reward delivery.

**Figure S3:**
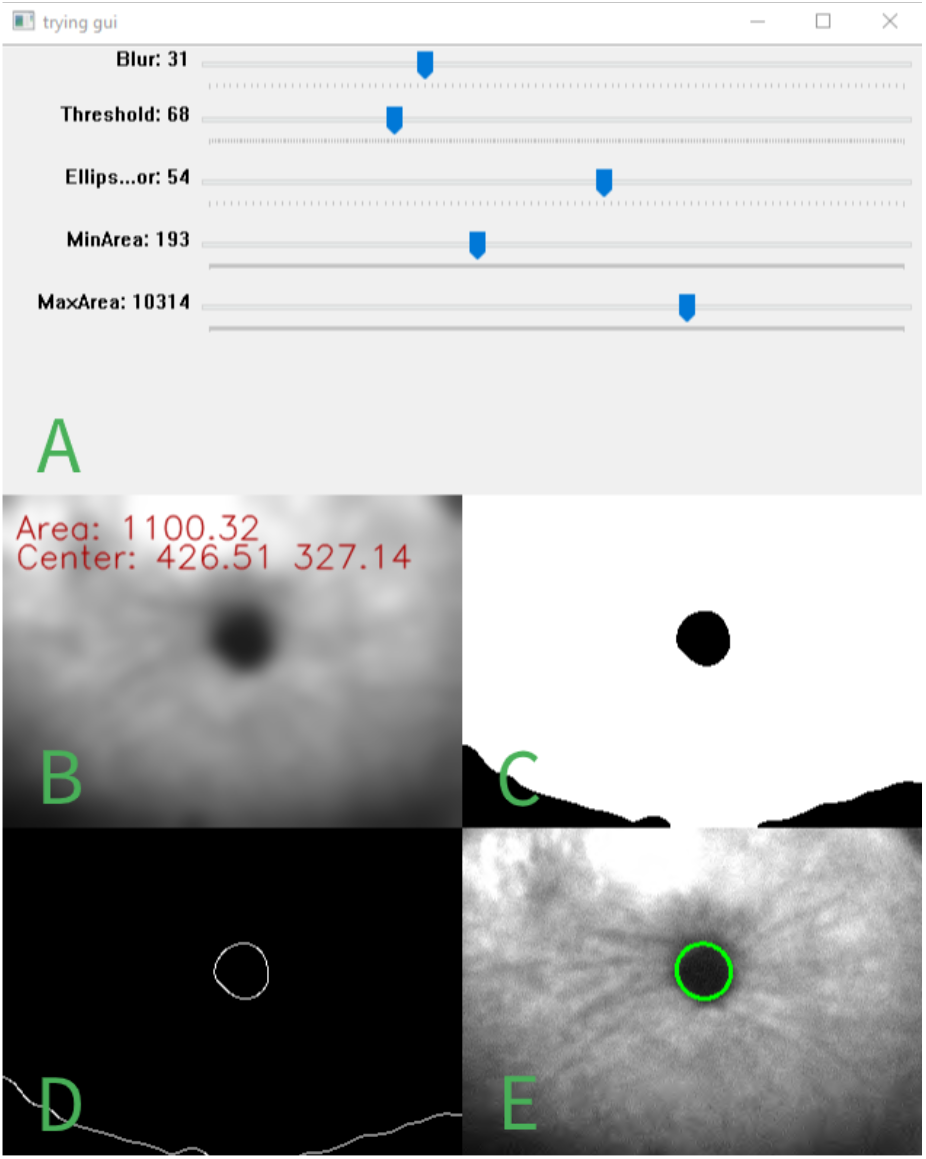
An image showing the GUI used to perform pupillometry. This GUI was used to manually set parameters for pupil detection on a single, randomly selected video frame. Those parameters were then applied to the entire recording in an automated procedure. **(A)** The parameter values. **(B)** The original image subjected to a Gaussian blur. **(C)** The binary image. **(D)** Closed contours were selected. **(E)** An ellipse was fit to the contour corresponding to the pupil.

